# Does habitat restoration disturb? A case study of a shallow urban water reservoir in western India using cladoceran zooplankton

**DOI:** 10.1101/2021.06.18.448979

**Authors:** Mihir R. Kulkarni, Sameer M. Padhye

## Abstract

Anthropogenic stressors, including restoration activities, can have ecosystem wide impacts, reflecting in various biotic components, particularly the basal levels in the trophic webs. Functional traits link taxonomic diversity to ecosystem function, thereby enabling a better ecological assessment of ecosystem health. We studied the effects of restoration activities on the community structure and functional diversity of freshwater cladoceran zooplankton in an urban water reservoir. Samples were taken in the early and late phases of the restoration work. Cladoceran species community and functional composition was significantly different between the two phases. There was a considerable reduction in taxonomic richness, functional richness and redundancy in the late phase. Taxonomic beta diversity between the two phases was explained more by nestedness. Habitat degradation due to haphazard restoration measures such as destruction of littoral zone and arbitrary desilting in addition to the dumping of untreated sewage could have contributed to the decrease in species and functional richness within the reservoir.

## INTRODUCTION

Anthropogenic effects on environment greatly alter the biological and functional diversity via modifications of communities (Loreau et al., 2001). The potential drivers of such changes can be identified by studying variations in biological communities and their environments and through integrative approaches using functional traits along with species identities (Cairns & Pratt, 1993, Nevalainen & Luoto, 2017). Functional traits include morphological, physiological and life history characters of organisms which directly link taxonomic diversity and ecosystem function (see Webb et al., 2010), and their integration in monitoring frameworks can vastly improve the understanding of the ecological impacts of anthropogenic disturbance (see Mason et al., 2005, Laliberte & Legendre, 2010, Stamou et al., 2019).

Freshwater ecosystems are highly impacted with human mediated alterations to their morphology, hydrology and nutrient biogeochemistry (Ormerod et al., 2010, Carpenter et al., 2011). These effects lead to community level changes in multiple organisms integral to trophic webs, thereby affecting ecological function and stability (Tallberg et al., 1999, Dodds, 2006) in natural and artificial habitats.

Urban water reservoirs are created primarily to provide water for agriculture and human use, but they also perform valuable roles as recreational areas, aquaculture spaces, biodiversity reserves and buffers against extreme weather events (e.g. rainfall) (Jurczak et al., 2019). Such reservoirs are subject to many stressors like extrinsic nutrient loading, high siltation and inorganic pollution thereby affecting the structure and function of aquatic communities at multiple levels (Hill et al., 2017). The resulting decline of ecosystem health thus prompts implementation of restoration measures i.e. efforts to mitigate and possibly reverse the negative effects caused by such anthropogenic disturbances.

Zooplankton are an important basal component of aquatic food webs and have been used for studying various ecological processes (Dumont & Negrea, 2002, Simões et al., 2015, Jurczak et al., 2019, Liu et al., 2020). Their crucial role in the ecosystem function, particularly their top-down control on phytoplankton (Stamou et al., 2019) gives them indicative value in assessing the ecosystem stability, trophic state as well as the success of restoration measures using functional traits (e.g. using body size) (Dussart & Defaye, 2001, Dumont & Negrea, 2002, Louette et al., 2008, Rizo et al., 2017).

We conducted this study to assess the effects of habitat modification (performed as part of restoration measures) on the zooplankton community in an urban reservoir in western India. This is especially important in the Indian context since habitat modification/destruction; insufficient sewage/effluent treatment and introduction of non-native species are pertinent problems leading to loss of aquatic species (Kharat et al., 2001, Reddy & Char, 2006). We used Cladocera (water fleas) as the zooplankton group owing to their diversity, microhabitat preferences, wide range of functional traits and basal trophic role (Dumont & Negrea, 2002). Specific questions addressed were, 1) how the habitat modifications that happened as a part of a restoration work altered the cladoceran community composition and 2) whether this affected the functional composition and diversity (suggestive of altered ecosystem functioning). We have also briefly commented on whether this restoration work resulted in BH of the zooplankton fauna in the reservoir.

## MATERIALS AND METHODS

### Study site

Pashan lake (18°31’59”N; 73°47’18”E) is situated to South West of the Pune City. The lake arose as the waters of the Ramnadi river, Pune were dammed ca. 1900’s for irrigation and domestic use. Since then, a rapid increase in residential area, industry and defence institutions has occurred along the lake. With decreasing water quality over the years due to high siltation, eutrophication etc., restoration was carried out in 2004-5 and 2010-11, followed by beautification work around the periphery of the lake in 2013. Under these activities de-silting, creation of an artificial island, re-contouring of the lake, construction of walls and stocking with non-native fish (*Oreochromis* sp.) etc. was carried out, amounting to large scale habitat modification (Fig. 1, Yardi et al., 2019, Author’s pers. obs.).

**Figure 1.**
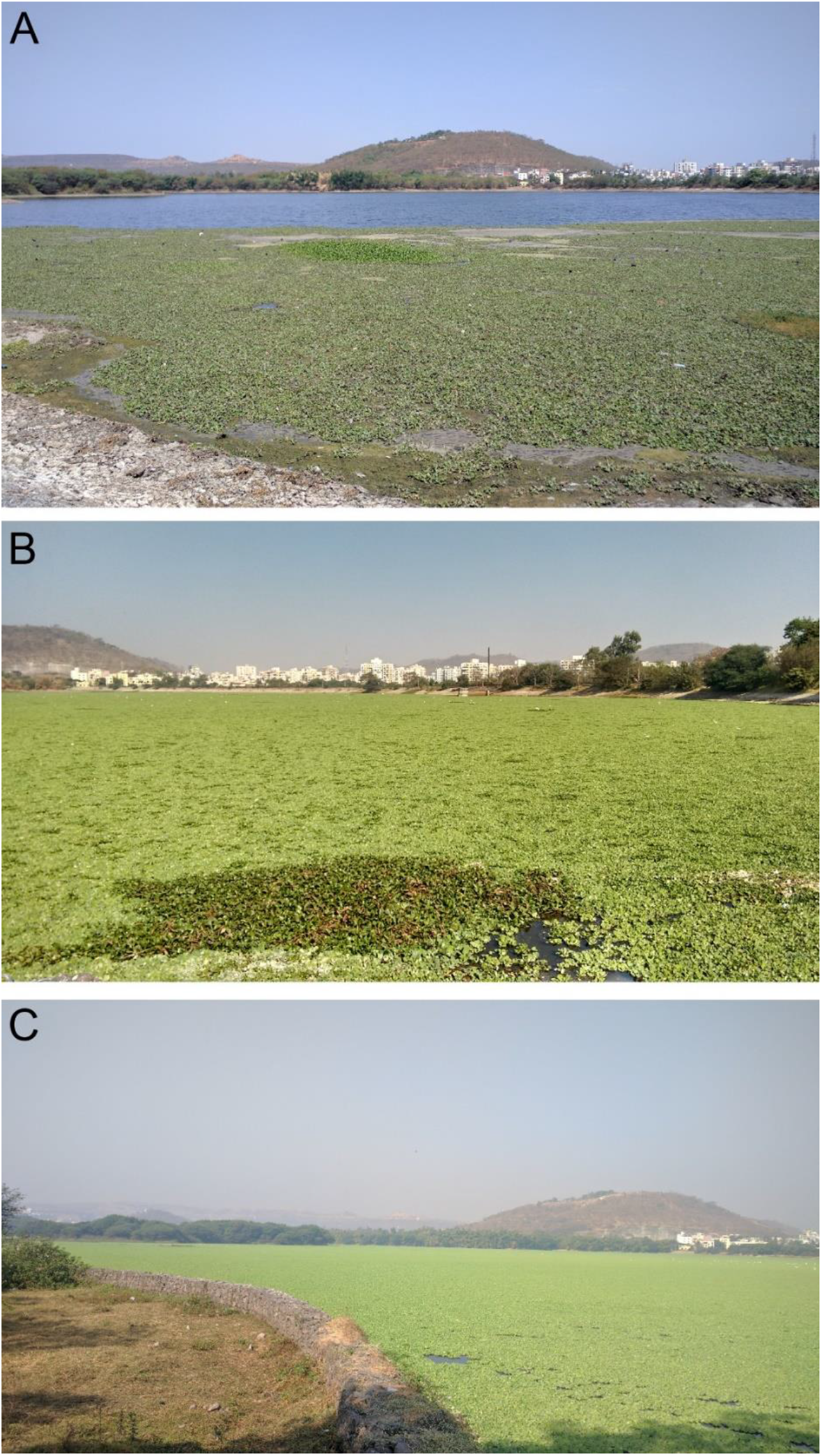
Photographs of Pashan reservoir taken from sampling stations showing the A, B) cover of floating macrophytes (predominantly *Eichornia* and *Pistia*) and C) habitat alterations – constructed retaining walls made of rocks wrapped in chain link.

Before this modification, submerged and emergent aquatic vegetation was commonly observed in the lake with species of *Vallisneria, Hydrilla, Ottelia, Typha, Chara* with floating *Pistia, Ipomoea* in low densities (SMP pers. obs.). A diverse avifauna was observed in the lake, with multiple migratory species being sighted thus making it a popular birding site. However, post-restoration and beautification, a decline in bird diversity associated with water quality was observed and reported. (Detailed information given here: https://www.indiawaterportal.org/articles/threatened-urbanisation-doomed-restoration).

### Sample collection

As a part of a larger survey of the Cladocera of Pune region, samples from Pashan reservoir were collected during 2009 -10 (n = 6), and in 2016 (n = 5) following the same protocol. Samples were categorized into “early” phase (2009-10, soon after restoration activity) and a “late” phase (2016, 5 years post-restoration) respectively.

Representative qualitative sampling (i.e. sampling in the littoral zones, sampling from the littoral sediments by gently disturbing the substratum and open patches adjacent to the littoral zones) was carried out during the initial collection of 2009-10 at 3 stations using tow-and hand-nets (mesh size 80µ) and pooled together as a single sample. Sampling was carried out from one side of the reservoir since only that face was accessible for sampling. Samples were preserved in 4% formalin. Similar regime was followed in the late phase (2016) as well for a comparative analysis. Efforts were made to collect from the same sites during the late phase but sometimes this was not possible owing to a very high density of *Eichornia* and *Pistia* (Fig.1A).

### Identification of cladoceran zooplankton

Specimens from the samples were examined under a stereomicroscope (Magnus MS24) and identified using a compound microscope (Olympus CH20i) following standard taxonomic keys (Supplementary information R1).

### Data Analyses

We carried out species richness estimations for both the collection periods using samples to check the efficacy of our sampling. Bootstrap index was used for the estimations and calculations were done using the r package ‘vegan’.

Non-metric multi-dimensional scaling (nMDS) based ordination using Jaccard similarity was used to visualize the shift in cladoceran community composition between the two phases. The model was run using 999 random starts and 2 dimensions (k).

Differences in the fauna of the two collection periods was analysed using one-way non-parametric permutational MANOVA (PERMANOVA, Anderson, 2005) using the Jaccard index with 5000 permutations using the ‘adonis’ function from vegan package. Homogeneity of dispersion was checked beforehand using ‘betadisper’ function in the vegan package (results in supplementary information table 1).

β-diversity partitioning procedure (Baselga, 2010) was used to study the extent of change in the community by analysing the relative contribution of turnover (βsim) and nestedness (βnes) and overall β-diversity (measured as Jaccard dissimilarity, βjac). This was performed using the βpart package using species presence/absence data from samples of both the phases (Baselga & Orme, 2012).

Functional composition and diversity was calculated using a set of 10 biological traits (adopted from Barnett et al., 2007, Rizo et al., 2017) which included body size, egg clutch size, Ratio of eye size to total length, feeding type based on the type of filtration mechanism and habitat type (Supplementary information table 2 for details). Four traits were continuous and the remaining six were categorical. Functional composition of the community was visualized using the species occurrence and functional trait data with an average linkage hierarchical dendrogram in the FD package. Data on functional traits were used to calculate functional richness (hypervolume of species community in a trait hyperspace; ‘Fric’ henceforth) and functional redundancy (degree of surplus traits available in the community; ‘Fred’ henceforth) (Laliberté & Legendre, 2010) using the packages FD and SYNCSA (Debastiani & Pillar, 2012). Significance of variation in the functional diversity components between the two collection periods was evaluated using the Wilcoxon signed rank test.

All analyses were performed using RStudio (v.3.6.2)

## RESULTS

### Altered community composition

The early phase had higher species richness per sample than the late phase (Fig.2A), comprising of 28 species from 7 families and 9 species from 4 families, respectively. Species such as *Pseudochydorus bopingi* Sinev et al. 2016 and *Oxyurella singalensis* (Daday, 1862) were restricted to early phase while species like *Moina micrura* Kurz, 1874 were observed in both the phases (Table 1). Chydoridae was the most species rich family in both the collection phases while certain families (e.g. Ilyocryptidae) were absent in the late phase (Fig.2B). Our sampling captured 80% of the estimated fauna for both early as well as late phase (Estimated species in the early and late phases = 32 and 11, respectively).

**Figure 2.**
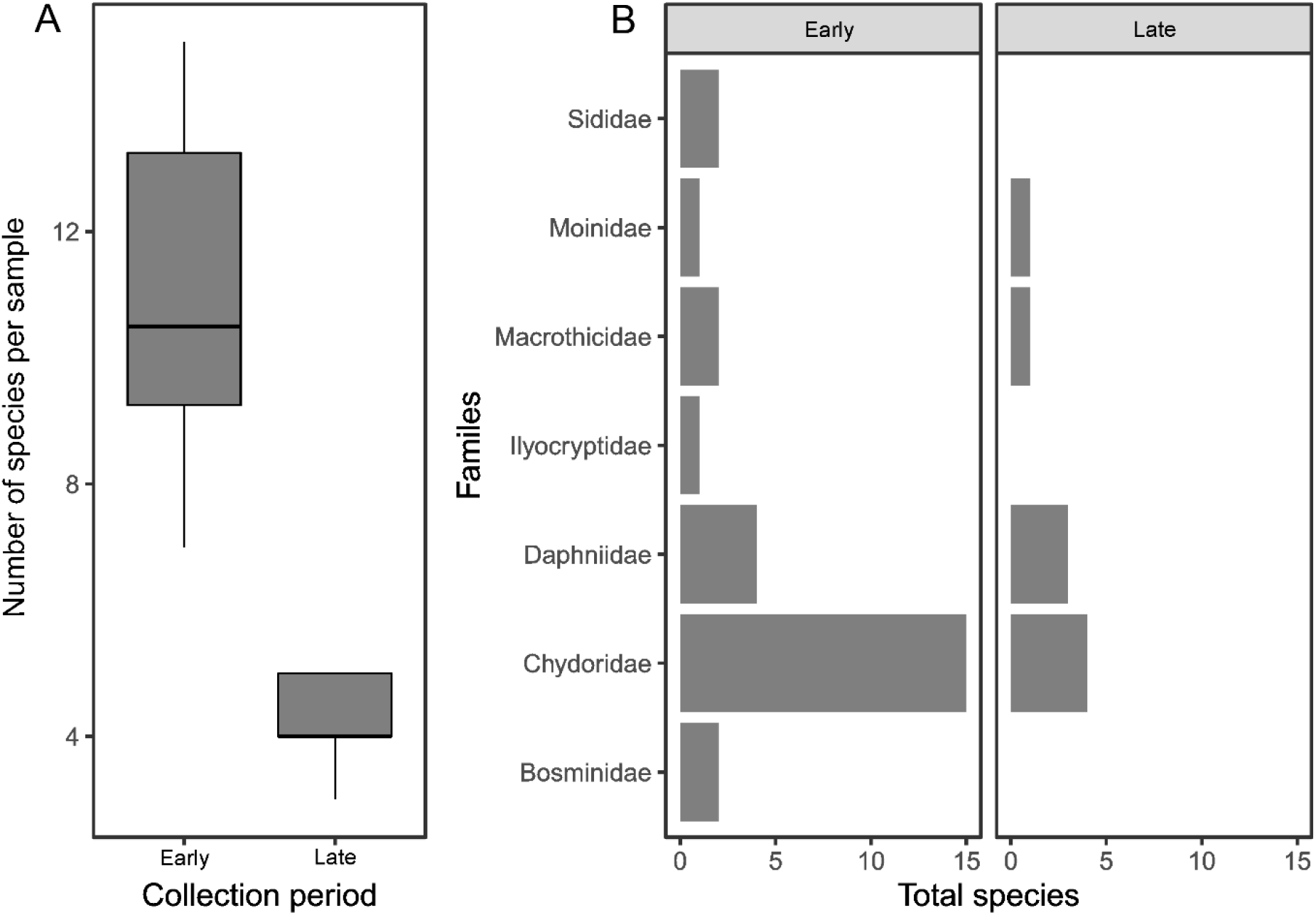
Reduction in cladoceran diversity over the study period observed in A) number of species per sample and B) family-wise composition.

**Table 1.**
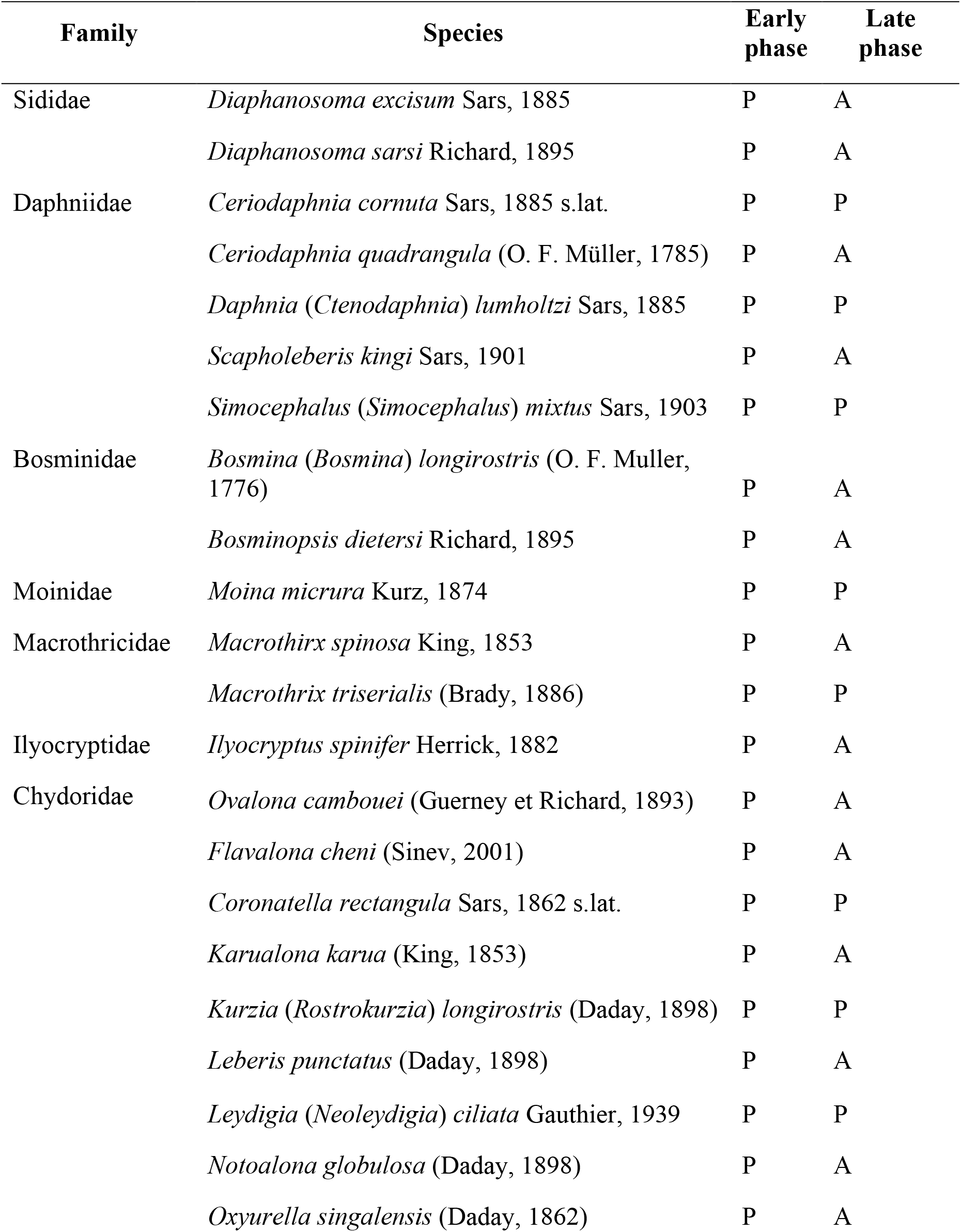
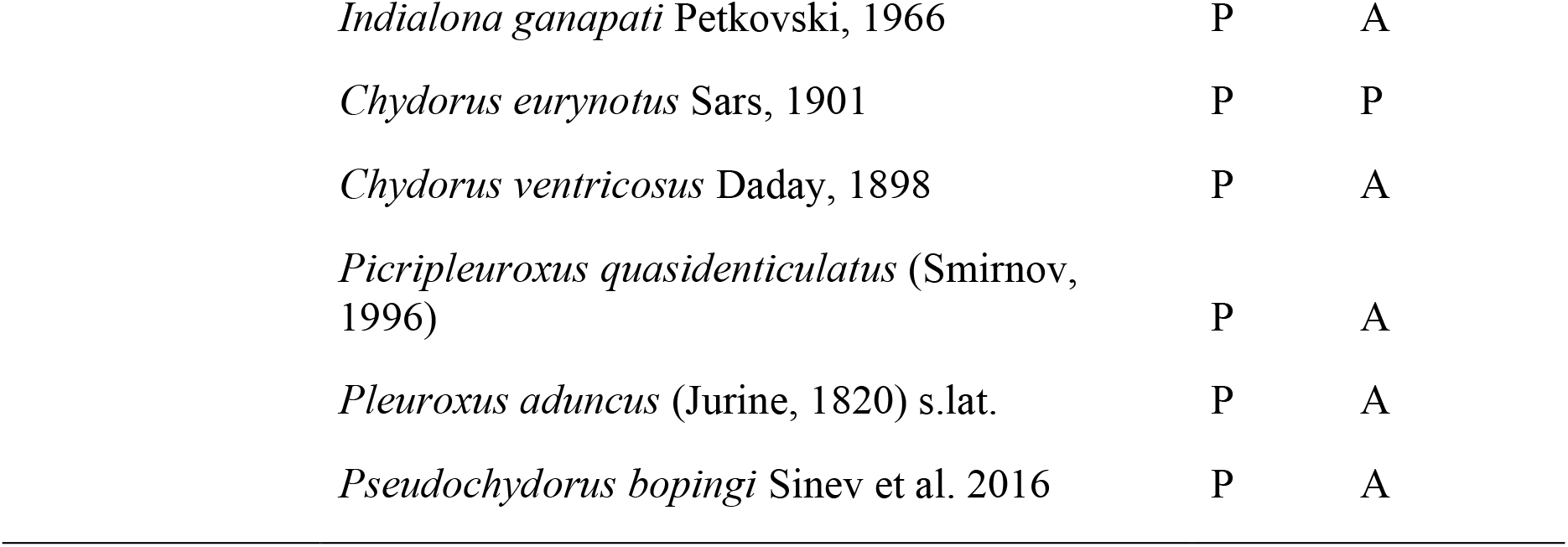
List of observed cladoceran species at Pashan in early and late phases of restoration (P: Present; A: Absent).

| Family | Species | Early phase | Late phase |
| --- | --- | --- | --- |
| Sididae | *Diaphanosoma excisum* Sars, 1885 | P | A |
|  | *Diaphanosoma sarsi* Richard, 1895 | P | A |
| Daphniidae | *Ceriodaphnia cornuta* Sars, 1885 s.lat. | P | P |
|  | *Ceriodaphnia quadrangula* (O. F. Müller, 1785) | P | A |
|  | *Daphnia* (*Ctenodaphnia*) *lumholtzi* Sars, 1885 | P | P |
|  | *Scapholeberis kingi* Sars, 1901 | P | A |
|  | *Simocephalus* (*Simocephalus*) *mixtus* Sars, 1903 | P | P |
| Bosminidae | *Bosmina* (*Bosmina*) *longirostris* (O. F. Muller, 1776) | P | A |
|  | *Bosminopsis dietersi* Richard, 1895 | P | A |
| Moinidae | *Moina micrura* Kurz, 1874 | P | P |
| Macrothricidae | *Macrothrix spinosa* King, 1853 | P | A |
|  | *Macrothrix triserialis* (Brady, 1886) | P | P |
| Ilyocryptidae | *Ilyocryptus spinifer* Herrick, 1882 | P | A |
| Chydoridae | *Ovalona cambouei* (Guerney et Richard, 1893) | P | A |
|  | *Flavalona cheni* (Sinev, 2001) | P | A |
|  | *Coronatella rectangula* Sars, 1862 s.lat. | P | P |
|  | *Karualona karua* (King, 1853) | P | A |
|  | *Kurzia* (*Rostrokurzia*) *longirostris* (Daday, 1898) | P | P |
|  | *Leberis punctatus* (Daday, 1898) | P | A |
|  | *Leydigia* (*Neoleydigia*) *ciliata* Gauthier, 1939 | P | P |
|  | *Notoalona globulosa* (Daday, 1898) | P | A |
|  | *Oxyurella singalensis* (Daday, 1862) | P | A |

|  |  |  |
| --- | --- | --- |
| *Indialona ganapati* Petkovski, 1966 | P | A |
| *Chydorus eurynotus* Sars, 1901 | P | P |
| *Chydorus ventricosus* Daday, 1898 | P | A |
| *Picripleuroxus quasidenticulatus* (Smirnov, 1996) | P | A |
| *Pleuroxus aduncus* (Jurine, 1820) s.lat. | P | A |
| *Pseudochydorus bopingi* Sinev et al. 2016 | P | A |

The cladoceran community composition was significantly different in the early and late phases, visualized as two distinct clusters (nMDS stress= 0.08) (Fig.3; Table 2A). The overall beta diversity was high between the early and late phases, and entirely driven by nestedness (βjac = βnes = 0.66; βsim = 0).

**Figure 3.**
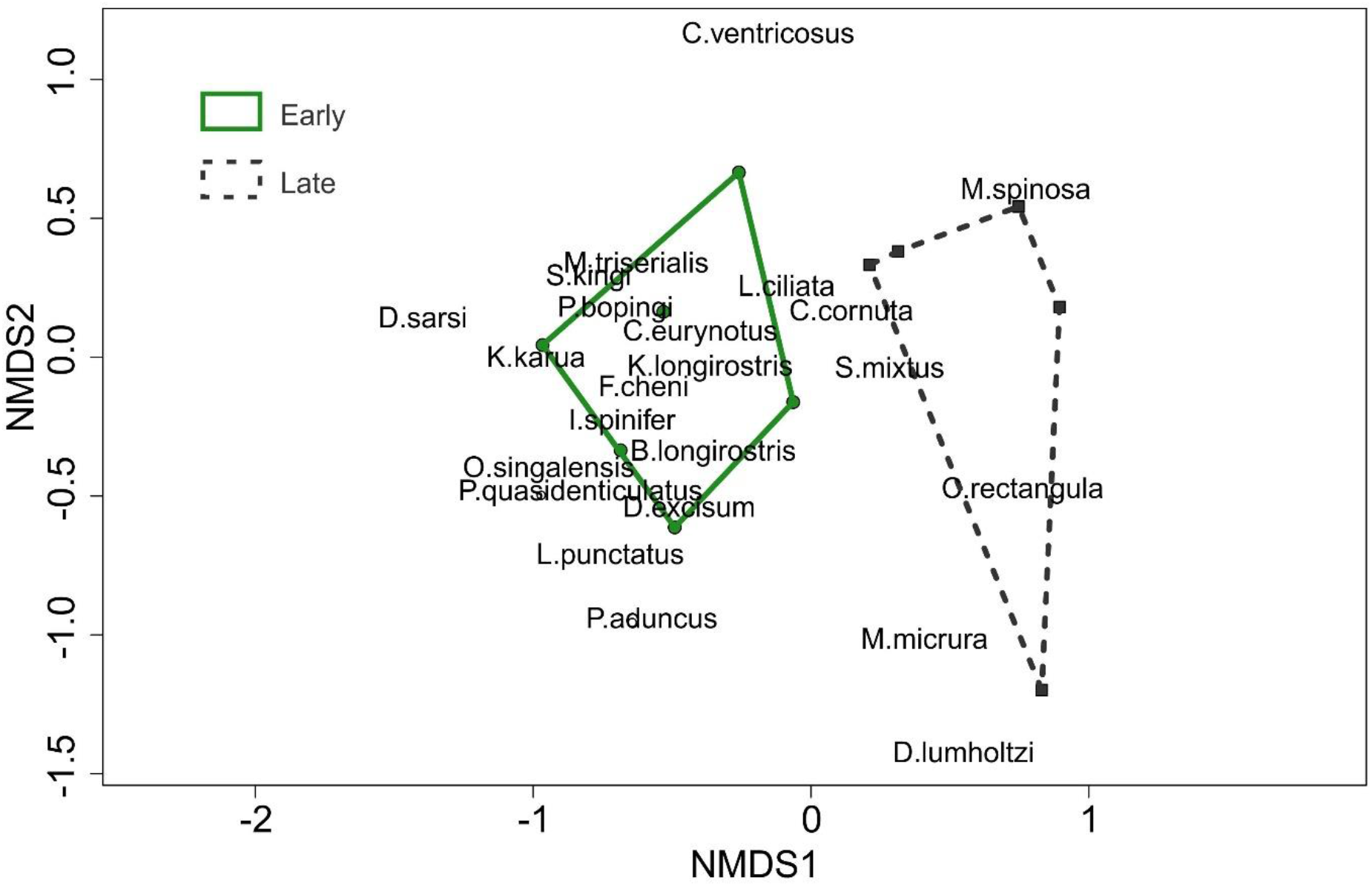
nMDS ordination showing clear separation among the early and late Cladocera communities at the study site (short form of species names used).

**Table 2.**
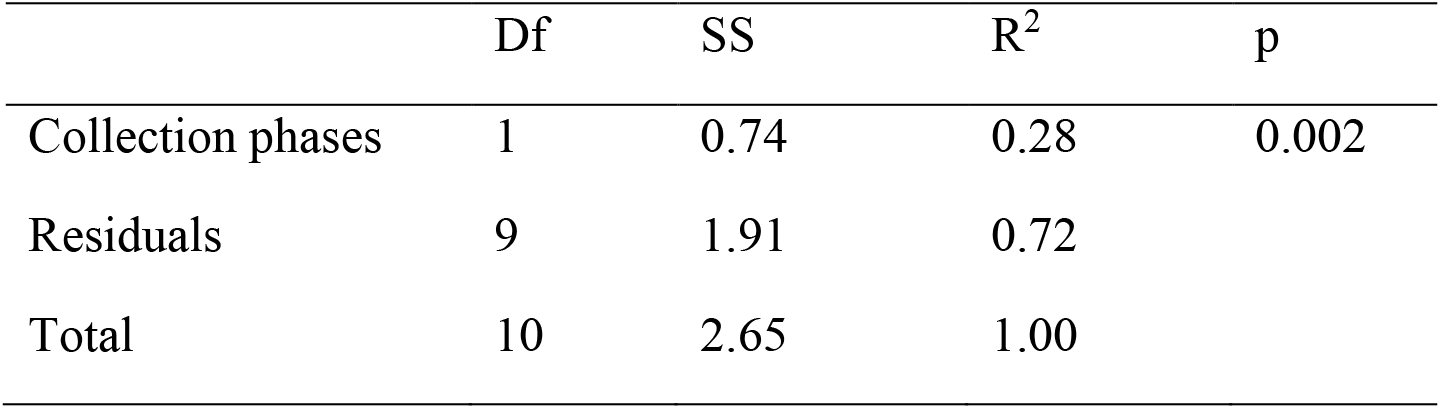
Results of one-way PERMANOVA for assessing the significance of differences in species community composition.

|  | Df | SS | R<sup>2</sup> | p |
| --- | --- | --- | --- | --- |
| Collection phases | 1 | 0.74 | 0.28 | 0.002 |
| Residuals | 9 | 1.91 | 0.72 |  |
| Total | 10 | 2.65 | 1.00 |  |

### Changes in functional composition

The functionally diverse community in the early phase shifted to a functionally poorer community in the late phase (Fig.4). The habitat preferences and food filtration types of observed species primarily resolved the functional groups. The maximum number of functional groups characterized reduced from six (range 2-6; F1-6 henceforth) in the early phase to three (range 2-3; F2,3,6) in the late phase (Fig.4). Of the total species, > 50 % were represented by functional group 3 comprising of small sized scrapers (Supplementary Information table 3 for details on each group) in both the phases. The functional composition of the late phase revealed presence of daphniid littoral filter feeders, pelagic feeders, a few littoral scrapers with the loss of bacteria filtering feeders, benthic pickers and non-daphniid littoral filter feeders which were observed in the early phase.

**Figure 4.**
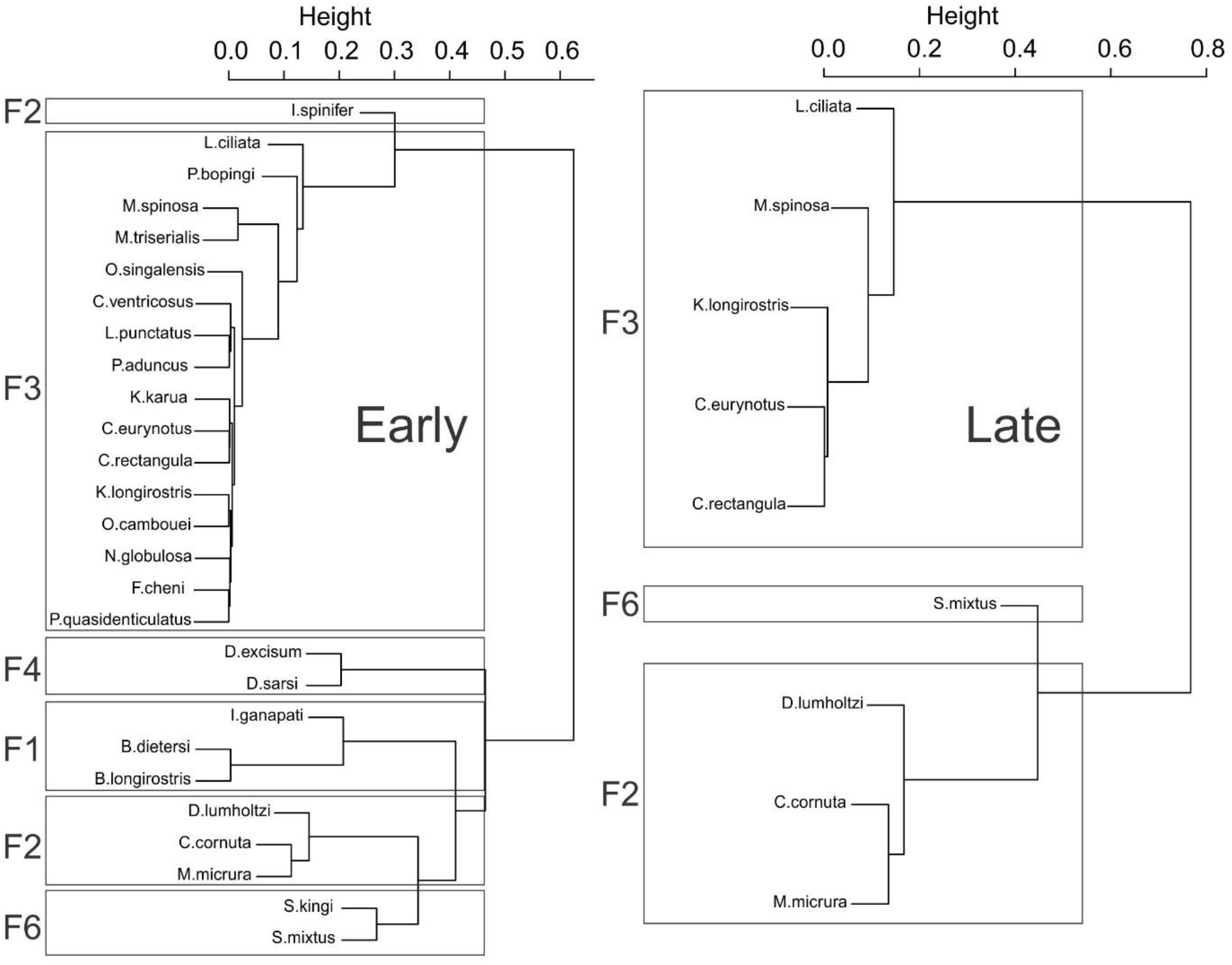
Cluster dendrograms of the functional composition of cladoceran communities in A) early and B) late phases of restoration. Each coloured box represents a separate functional group (For characteristics of each group, please refer to supplementary information table 2 & table 3 for details on functional groups F1-6).

### Changes in functional richness and redundancy

Values of functional richness and redundancy were lower in the late phase (Fig. 5; Fric: early mean: 0.2 ± 0.06 s.d.; late mean: 0.04 ± 0.02 s.d.; Fred: early mean: 0.37 ± 0.05 s.d.; late mean: 0.27 ± 0.04 s.d.) and significantly different between the two periods (Fric: W = 30; p = 0.04; Fred: W = 30; p = 0.043) (For all values, refer to Supplementary Information table 4).

**Figure 5.**
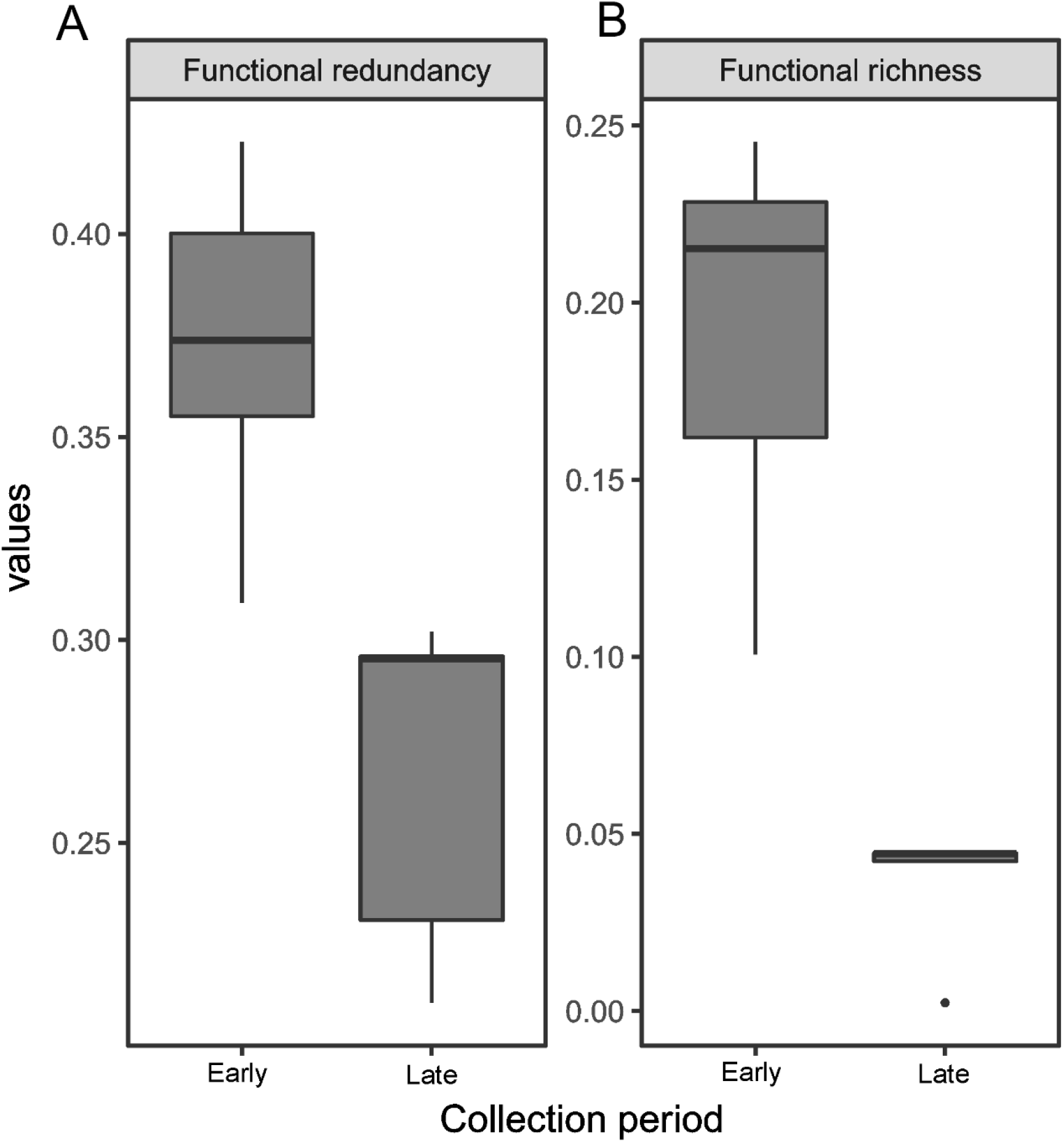
Reduction in A) functional redundancy and B) functional richness of the cladoceran community over the early and late phases.

## DISCUSSION

Our study on the zooplankton community of a eutrophic urban reservoir specifically revealed a) species richness decreased with only a small subset of the early community observed during the late phase, b) functional composition became species poorer and functional redundancy reduced in the late phase.

Zooplankton species richness could be influenced by structural changes to lake topography, which can alter specific zones of the ecosystem. The littoral zone, characterised by lower depth, high light penetration and presence of aquatic macrophytes is important for many taxa like the zooplankton molluscs, fish, amphibians, and waterbirds (Thorp & Covich, 2001, Santos et al., 2018). Littoral macrophytes comprise of submerged and emergent species, which increase the overall habitat complexity by creating specific micro-habitats for various species including zooplankton (DiFonzo et al., 1988, Patterson, 1993, Van Onsem et al., 2010) and birds (especially waders and shorebirds). Littoral vegetation commonly observed in the lake in the early phase (*Vallisneria, Hydrilla, Ottelia, Typha, Chara* and floating *Pistia, Ipomoea* in low densities; SMP pers. obs.) was completely absent in the late phase and replaced entirely by *Ipomea* and *Pistia* sp. (Fig.1A-C).

Removal of bottom sediment from reservoirs is widespread (Reddy & Char, 2006, Jurczak et al., 2019). It can directly influence zooplankton communities by affecting the local egg bank structure (Maia-Barbosa et al., 2003, Wojtal-Frankiewicz et al., 2019). These “egg banks” are crucial for resurrection of the community after dry phases. Although egg banks have large number of eggs with a long viability, most “active” stages occur in the top layers (Brendonck & De Meester, 2003), which are prone to mechanical removal (for desilting/reducing internal phosphate loading) or capping to reduce internal phosphate loading during restoration measures. As a result, these activities can lead to reduction in the “native” community. Such changes in the bathymetric profile (due to desilting) in combination with a shift in aquatic vegetation could have led to loss of associated zooplankton species (particularly littoral scrapers) from the reservoir (Table 1, Fig. 4).

Pashan restoration work clearly resulted in physical modification of the habitat with loss of microhabitats. The decrease in species richness that occurred due to this disturbance event and the high nestedness observed amongst the early-and late-communities indicate the effect of a hierarchical species loss (Baselga, 2010). Similar patterns of high nestedness in zooplankton communities were observed in disturbed habitats (Gianuca et al., 2017, Braghin et al., 2018). Functional diversity of communities provides information on microhabitats, resource use and trophic dynamics in a habitat (Jeppesen et al., 2000), as it connects taxonomic diversity to ecosystem function (Nevalainen & Luoto, 2017). Large shifts in environment can change functional composition with replacement and/or substitution of functional traits (Loreau et al., 2001, Petchey & Gaston, 2002, Slade et al., 2007). This was evident as the loss of functional groups and decrease in the functional richness in the late phase of the study (Fig.4), observed in the zooplankton community. The late phase cladoceran community predominantly comprised of smaller sized species (Ex. *Ceriodaphnia cornuta*). A higher functional redundancy reflects available ecological alternatives to some taxa in the habitat, thus ensuring maintenance of ecosystem function in case of loss of taxonomic diversity to a certain extent (Walker, 1992, Naeem, 1998). However, this also underscores the crucial role of functional redundancy in ecosystem resilience to disturbances (Rosenfeld, 2002). Although three of the cladoceran functional groups (F2, 3, 6) observed in the early community were retained, there was a significant reduction in number of species from those groups thus affecting the redundancy and highlighting the disturbance of ecosystem function and its vulnerability to further disturbance (Fig.4).

Increased nutrient loading commonly affects urban reservoirs, and remains one of the major drivers of eutrophication. Recent reports showed a relatively high phosphate loading (approx. 0.49 mg/l) in Pashan lake which could promote rapid growth of algae and floating macrophytes (Fig.1) (Smith et al., 1999, Smith, 2003, Yardi et al., 2019) thereby affecting the zooplankton communities (Nevalainen & Luoto, 2017). We were not able to measure the environmental variables due to logistical reasons and thus refrain from making any conjectural statements. The biotic communities in urban aquatic habitats are often “homogenized” due to multiple stressors (McKinney & Lockwood, 1999, Rahel, 2002, Liu et al., 2020, Lougheed et al., 2008). Biotic homogenization can occur via displacement of rare or endemic species with widespread ones (McKinney & Lockwood, 1996) and major structural changes in communities due to reduced taxonomic and/or functional diversity, all of which was observed during the late phase of the reservoir. Faunal homogenization in urban aquatic habitats has been reported in several taxa including zooplankton (Liu et al., 2020, Devictor et al., 2008, Ibáñez-Álamo et al., 2017). Cladoceran species found in the late phase mostly consisted of many regionally common species such as *Ceriodaphnia cornuta, Simocephalus mixtus* and *Moina micrura* (for categorization of cladoceran species occurrences, please refer to Padhye & Dumont (2014)). Data from Yardi et al. (2019) points to an avifauna of Pashan lake composed largely (>50%) of regionally abundant species, similar to those reported by Koparde and Raote (2016). These observations suggest biotic homogenization occurring at multiple levels, and require additional focussed studies to identify the drivers, which is crucial for further management.

## ACKNOWLEDGEMENTS

We thank International Society of Limnology (SIL), Tonolli award, Council for Scientific and Industrial Research, India (CSIR) research fellowship (0556/2014/EMR1) for providing the funding. We also acknowledge Dr. Shruti Paripatyadar (Pune) for help with sampling and discussions, Dr. Hemant Ghate (Modern College, Pune) for helpful discussions. Dr. Kalpana Pai (Dept. of Zoology, SPPU, Pune) and Director (Biologia Life Science LLP, Ahmednagar, India) for their support.

