## Supplementary information for "Does habitat restoration disturb? A case study of a shallow urban water reservoir in western India using cladoceran zooplankton"

Table 1. Beta dispersion test results for PERMANOVA using taxonomic species data

|  | df | SS | F value | p |
| --- | --- | --- | --- | --- |
| Groups | 1 | 0.044 | 0.18 | 0.68 |
| Residuals | 9 | 0.22 |  |  |

Table 2. The traits used in calculation of functional diversity indices (for more details, please refer to Table 1 from Rizo et al. 2017)

| **Trait name** | **Type** | **Units/Categories** |
| --- | --- | --- |
| Average body size | Numeric-continuous | mm |
| Average Egg clutch size | Numeric- integer | number of eggs |
| Proportion of eye size to total length | Numeric-continuous | mm |
| Filtration type | Categorical | D-type/S-type/I-type/B-type/C-type |
| Habitat preference | Categorical | Pelagic/Littoral |
| Carapace colour | Categorical | Colored/Transparent |
| Proportion of eye size to total body size | Numerical | Percent (%) |
| Preferred trophic condition | Categorical | Oligo-Mesotrophic,  Meso-Eutrophic,  Eutrophic |
| Trophic Regime | Categorical | Herbivore,  Herb-Detritivore |
| Predator escape response | Categorical | pausing and jumping, rapid  swimming, not moving |

Table 3. Functional groups of cladocerans observed at Pashan.

| **Functional group** | **Name of the group** | **Diagnostic characters** |
| --- | --- | --- |
| F1 | Bacteria filtering filter feeders | Very small planktonic species having smaller clutch size using filtering for gathering very small sized feed. |
| F2 | Pelagic filter feeders | Large sized planktonic species having higher egg clutches and using 3rd and 4th trunk limbs for filtering food |
| F3 | Substratum scrapers | Relatively small sized littoral species having smaller clutch size and using scrapers on their second trunk limb for food gathering |
| F4 | Non Daphniid Littoral Filter feeders | Moderate sized littoral species having moderate egg clutch size using 1st - 5th trunk limbs for filtering food |
| F5 | Benthic pickers | Moderately sized benthic species having moderate number of eggs using special ‘picking’ mechanism to gather food from benthic sediments |
| F6 | Daphniid Littoral Filter feeders | Moderate to big sized species having higher egg clutches, preferring littoral zones and using 3rd and 4th trunk limbs for filtering food |

Table 4. Functional richness and redundancy values for all samples.

| **Collection group** | **Functional richness (Fric)** | **Functional redundancy (Fred)** |
| --- | --- | --- |
| Early | 0.100724 | 0.391519 |
| Early | 0.206853 | 0.309148 |
| Early | 0.147078 | 0.356126 |
| Early | 0.245345 | 0.403021 |
| Early | 0.223844 | 0.422622 |
| Early | 0.229944 | 0.354762 |
| Late | 0.002342 | 0.210632 |
| Late | 0.044623 | 0.295411 |
| Late | 0.042297 | 0.302033 |
| Late | 0.044477 | 0.295985 |
| Late | 0.044685 | 0.230994 |

R1. Additional references used for identifying Cladocera

Benzie, J.A.H. 2005. The genus Daphnia (including Daphniopsis) (Anomopoda: Daphniidae). Guides to the identification of the microinvertebrates of the continental waters of the world H.J.Dumont ed., SPB Academic Publishing: 1-383.

Berner, D.B. (1985) Morphological differentiation among species in the Ceriodaphnia cornuta complex (Crustacea,Cladocera). Verhandlungen der Internationalen Vereinigung fuer Theoretische und Angewandte Limnologie, 22, 3099–3103.

Dumont, H. J. & Pensaert, J. (1983) A revision of the Scapholeberinae (Crustacea: Cladocera). Hydrobiologia, 100, 3–45.http://dx.doi.org/10.1007/BF00027420

Dumont, H.J. & Silva–Briano, M. (2000) Karualona n.gen. (Anomopoda: Chydoridae), with a description of two new species, and a key to all known species. Hydrobiologia, 435, 61–82.

Dumont, H.J., Silva–Briano, M. & Subash Babu, K.K. (2002) A re-evaluation of the Macrothrix rosea–triserialis group, with the description of two new species (Crustacea Anomopoda: Macrothricidae). Hydrobiologia, 467, 1–44.

Goulden, C.E. (1968) The systematics and evolution of the Moinidae. Transactions of the American Philosophical Society Held at Philadelphia, new series, 58, 1–101. <http://dx.doi.org/10.2307/1006102>

Hudec, I, 1991. A comparison of populations from the Daphnia similis group (Cladocera: Daphniidae). Hydrobiologia 225: 9-22.

Hudec, I. (2000) Subgeneric differentiation within Kurzia (Crustacea: Anomopoda: Chydoridae) and a new species from Central America. Hydrobiologia, 421, 165–178.

Korovchinsky, N.M. (1992) Sididae & Holopediidae (Crustacea: Daphniiformes). Guides to the identification of the microinvertebrates of the continental waters of the world 3. SPB Academic Publishing, The Hague, 82 pp.

Kotov, A.A. 2000. Re-description and assignment of the chydorid Indialona ganapati Petkovski, 1966 (Branchiopoda: Anomopoda: Aloninae) to Indialonini, new tribus. Hydrobiologia 439: 161-178.

Kotov, A.A. (2009) A revision of Leydigia Kurz, 1875 (Anomopoda, Cladocera, Branchiopoda), and subgeneric differentiation within the genus. Zootaxa, 2082, 1–68.

Kotov, A.A. & Štifter, P. 2006. Ilyocryptidae of the world. Guides to the identification of the microinvertebrates of the continental waters of the world. Dumont, H.J., SPB Academic Publishing: 1-172.

Kotov, A.A., Ishida, S. & Taylor, D.J. (2009) Revision of the genus Bosmina Baird, 1845 (Cladocera: Bosminidae), based on evidence from male morphological characters and molecular phylogenies Zoological Journal of the Linnean Society, 156, 1–51.

Michael, R.G. & Sharma, B.K. (1988) Fauna of India and ajancent countries. Indian Cladocera (Crustacea: Branchiopoda: Cladocera). Zoological Survey of India, Calcutta, 262 pp.

Rajapaksa, R. & Fernando, C.H. (1986a) A review of the systematics and distribution of Chydorus ventricosus Daday, 1898, with the first description of the male and redescription of the species. Canadian Journal of Zoology, 64, 818–832. <http://dx.doi.org/10.1139/z86-123>

Rajapaksa, R. & Fernando, C.H. (1987c) Redescription and assignment of Alona globulosa Daday, 1898 to a new genus Notoalona and a description of Notoalona freyi sp. nov. Hydrobiologia, 144, 131–153. <http://dx.doi.org/10.1007/BF00014527>

Sinev, A.Y. (1999) Alona costata Sars, 1862 versus related palaeotropical species: the first example of close relations between species with a different number of main head pores among Chydoridae (Crustacea: Anomopoda). Arhropoda Selecta, 8(3), 131–148.

Sinev, A.Y. (2001c) Separation of Alona cambouei Guerne & Richard, 1893 from Alona pulchella King, 1853 (Branchiopoda: Anomopoda: Chydoridae). Arthropoda Selecta, 10(1), 5–18.

Sinev, A.Y., Van Damme, K. & Kotov, A.A. (2005) Redescription of tropical–temperate cladocerans Alona diaphana King, 1853 and Alona davidi Richard, 1895 and their translocation to Leberis Smirnov, 1989 (Branchiopoda: Anomopoda: Chydoridae). Arthropoda Selecta, 14(3), 183–205.

Sinev, A.Y. 2011. Re-description of the rheophilous Cladocera Camptocercus vietnamensis Than, 1980 (Cladocera: Anomopoda: Chydoridae). Zootaxa 2934: 53–60.

Sinev, A. Y., Garibian, P. G. & Gu, Y. 2016. A new species of Pseudochydorus Fryer, 1968 (Cladocera: Anomopoda: Chydoridae) from South-East Asia. Zootaxa 4079: 129–139.

Smirnov, N.N. 1971. Chydoridae fauny mira. Fauna USSR. Rakoobraznie, 1. Leningrad [English translation: Chydoridae of the world. Israel Program for Scientific Translations, Jerusalem, 1974]

Smirnov, N.N. 1992. The Macrothricidae of the world. Guides to the identification of the microinvertebrates of the continental waters of the world. Dumont, H.J., SPB Academic Publications: 1-143.

Smirnov, N.N. 1996. Cladocera: The Chydorinae and Sayciinae (Chydoridae) of the world. Guides to the identification of the microinvertebrates of the continental waters of the world. Dumont, H.J., SPB Academic Publications: 1-197.

Van Damme, K. & Dumont, H.J. 2008. The ‘true’ genus Alona Baird, 1843 (Crustacea: Cladocera: Anomopoda): characters of the A. quadrangularis group and description of a new species from Democratic Republic Congo. Zootaxa 1945: 1–25

Van Damme, K., Sinev, A.Y. & Dumont, H.G. 2011. Separation of Anthalona gen.n. from Alona Baird, 1843 (Branchiopoda: Cladocera: Anomopoda): morphology and evolution of scraping stenothermic alonines. Zootaxa 2875: 1–64.
